# Pupil-linked arousal biases evidence accumulation towards desirable percepts during perceptual decision-making

**DOI:** 10.1101/2020.05.29.124115

**Authors:** Yuan Chang Leong, Roma Dziembaj, Mark D’Esposito

**Affiliations:** Helen Wills Neuroscience Institute, University of California, Berkeley; Department of Management Science & Engineering, Stanford University; Department of Psychology, University of California, Berkeley

## Abstract

People are biased towards seeing outcomes they are motivated to see. The arousal system coordinates the body’s response to motivationally significant events, and is well positioned to regulate motivational effects on sensory perception. However, it remains unclear whether arousal would enhance or reduce motivational biases. Here we measured pupil dilation as a measure of arousal while participants performed a visual categorization task. We used monetary bonuses to motivate participants to see one category over another. Even though the reward-maximizing strategy was to perform the task accurately, participants were more likely to report seeing the motivationally desirable category. Furthermore, higher arousal levels were associated with making motivationally biased responses. Analyses using computational models suggest that arousal enhanced motivational effects by biasing evidence accumulation in favor of motivationally desirable percepts. These results suggest heightened arousal biases people towards what they want to see and away from an objective representation of the environment.

**Statement of Relevance:** When confronted with an event of motivational significance (e.g., an opportunity to earn a huge reward), people often experience a strong arousal response that includes increased sweating, faster heart-rate and larger pupils. Does this arousal response help individuals make more accurate decisions, or does it instead bias and impair decision-making? This work examines the effects of arousal on how people decide what they see when they are motivated to see a particular outcome. We found that heightened arousal, as measured by larger pupils, was associated with a bias in how participants accumulated sensory evidence to make their decisions. As a result, participants became more likely to report seeing an ambiguous visual image as the interpretation they were motivated to see. Our results suggest that arousal biases perceptual judgments towards desirable percepts, and that modulating arousal levels could be a promising approach in reducing motivational biases in decision-making.

---

Imagine playing a heated tennis match and hitting a shot that might or might not have grazed the sideline. Would your motivation to win the point make you more likely to see the ball as having stayed within bounds? For most real-world perceptual decisions, people are not neutral observers indifferent to different perceptual outcomes. Some outcomes are better than others, and people are *motivated* to see those percepts over alternatives. Evidence from a number of studies suggest that wanting to see a desirable outcome biases people towards seeing that outcome, a phenomenon known as motivated perception (Balcetis et al., 2012; Balcetis & Dunning, 2006; Leong et al., 2019; Voss et al., 2008). For example, when presented with a visually ambiguous line drawing, participants were more likely to report seeing the percept associated with a desirable outcome (Balcetis & Dunning, 2006). Motivated perception has been shown to impair perceptual decision-making by biasing people towards what they want to see and away from the objective representation of external stimuli (Leong et al., 2019). Although previous studies provide growing evidence that motivation influences perception, it is not yet known how the interaction between motivation and sensory processing occurs.

The arousal system is well positioned to mediate motivational influences on perceptual processes. The level of physiological arousal performs an important role in coordinating the body’s response to motivationally significant events, such as the opportunity to obtain potential rewards or the appearance of an imminent threat (Lang & Bradley, 2010). Motivationally relevant stimuli activate arousal circuits and trigger an autonomic nervous system response that includes changes in heart rate, pupil dilation and skin conductance. Fluctuations in arousal are thought to be regulated by the locus coeruleus norepinephrine system (Sara & Bouret, 2012), and have been shown to impact sensory processing, memory encoding and decision-making (Aston-Jones & Cohen, 2005; Markovic et al., 2014; Mather et al., 2016). Furthermore, recent studies have shown that arousal influences both the accumulation of sensory information and response biases during perceptual decision-making (de Gee et al., 2020; Keung et al., 2019; Krishnamurthy et al., 2017; Murphy et al., 2014; Urai et al., 2017). These past studies, however, do not examine the role of arousal in a context where participants are motivationally biased to see one percept over another. It is thus unclear if and how arousal modulates motivational biases in perceptual decision-making.

How might arousal be related to the processes underlying motivated perception? According to the “glutamate amplifies noradrenergic effects” (GANE) model (Mather et al., 2016), arousal-related norepinephrine release interacts with local glutamate levels to selectively amplify the neural representation of physically or emotionally salient stimuli (see also Markovic et al. 2014). Consistent with this account, salient stimuli are preferentially perceived and remembered under heightened arousal (Kensinger et al., 2007; Lee et al., 2018). Building on this past work, we hypothesize that otherwise neutral stimuli become *motivationally salient* when participants are motivated to see them, and that arousal selectively biases sensory processing in favor of these stimuli. As such, we predict that arousal would be associated with stronger motivational biases during perceptual decision-making.

In the current work, we combined psychophysics, computational modeling and pupillometry to examine the relationship between arousal and perceptual decision-making. Participants were presented with visually ambiguous images and were rewarded for correctly categorizing the image into one of two categories. Pupil diameter was recorded during the task as a proxy measure for physiological arousal (Bradley et al., 2008). In different experimental blocks, we motivated participants to see one category over another by instructing them they would win extra money if the block contained more images from one category. Using a drift diffusion model (Ratcliff & McKoon, 2007), we modeled participants’ responses as the stochastic accumulation of sensory evidence towards a decision threshold. We then assessed if pupil diameter was associated with either or both motivational biases in the starting point and rate of evidence accumulation. By combining physiological markers of arousal with computational modeling, our study provides a mechanistic account of how arousal and motivation interact to change the way we perceive the environment. Our results help refine existing theories on arousal, and provide new insights into the role of affective states in regulating human cognition.

## Methods

### Participants

Forty-one participants were recruited from the [institution masked for review] community (sample of convenience) for a targeted sample size of 36, which a power analysis with effect estimates obtained from a previous study (Leong et al., 2019) indicated had greater than 80% power to detect a difference in psychometric curves between conditions. All participants provided informed consent prior to the start of the study. Participants were paid between US$20 and $30 depending on their performance on the task. Data from three participants were excluded due to unsuccessful eye-tracker calibration, yielding an effective sample size of thirty-eight participants (15 male, 23 female, 18-40 years of age, mean age = 21 years). All experimental procedures were approved by the [institution masked for review] Committee for the Protection of Human Subjects.

### Stimuli

For each participant, 12 sets of composite face/scene stimuli were created. Each stimulus set consists of 25 grey-scale images comprising a mixture of a face image and a scene image in varying proportions. Pilot data (n = 30) indicated that participants were equally likely to categorize an image as face-dominant or scene-dominant when the face-scene proportion was 48% scene and 52% face (i.e. point of subjective equivalence; *Supplemental Material*, Fig. S1). Thus, images with less than 48% scene were considered face-dominant while images with more than 48% scene were considered scene-dominant. Half of the stimulus sets contained more scene-dominant images (1 × 33% scene, 3 × 43% scene, 16 × 48% scene, 3 × 53% scene, 2 × 63% scene), while the other half contained more face-dominant images (2 × 33% scene, 3 × 43% scene, 16 × 48% scene, 3 × 53% scene, 1 × 63% scene). All stimuli were created to be isoluminant. Face images were frontal photographs posing a neutral expression, and were taken from the Chicago Face Database (Ma et al., 2015). Stimuli were presented using MATLAB software and the Psychophysics Toolbox (Brainard, 1997).

### Experimental task

On each trial, participants were presented with a face-scene composite image and had to categorize whether the image was face-dominant or scene-dominant (Fig. 1A). The image was presented for 3s and participants had to respond during this time. Participants earned 5 cents for each correct categorization. During the inter-trial interval (ITI; range = 2-6 seconds, mean = 3.5 seconds), a scrambled image with the same average luminance was presented to minimize luminance change on screen. Participants performed 12 blocks of 25 trials. In four of the blocks, we motivated participants to see face-dominant images by instructing them that they would win a $3.00 bonus if the block had more face-dominant images (*Face Bonus blocks*). In another four blocks, we motivated them to see scene-dominant images by instructing them that they would win a $3.00 bonus if the block had more scene-dominant images. In the remaining four blocks, participants performed the task without a motivation manipulation (*Neutral blocks*).

**Figure 1.**
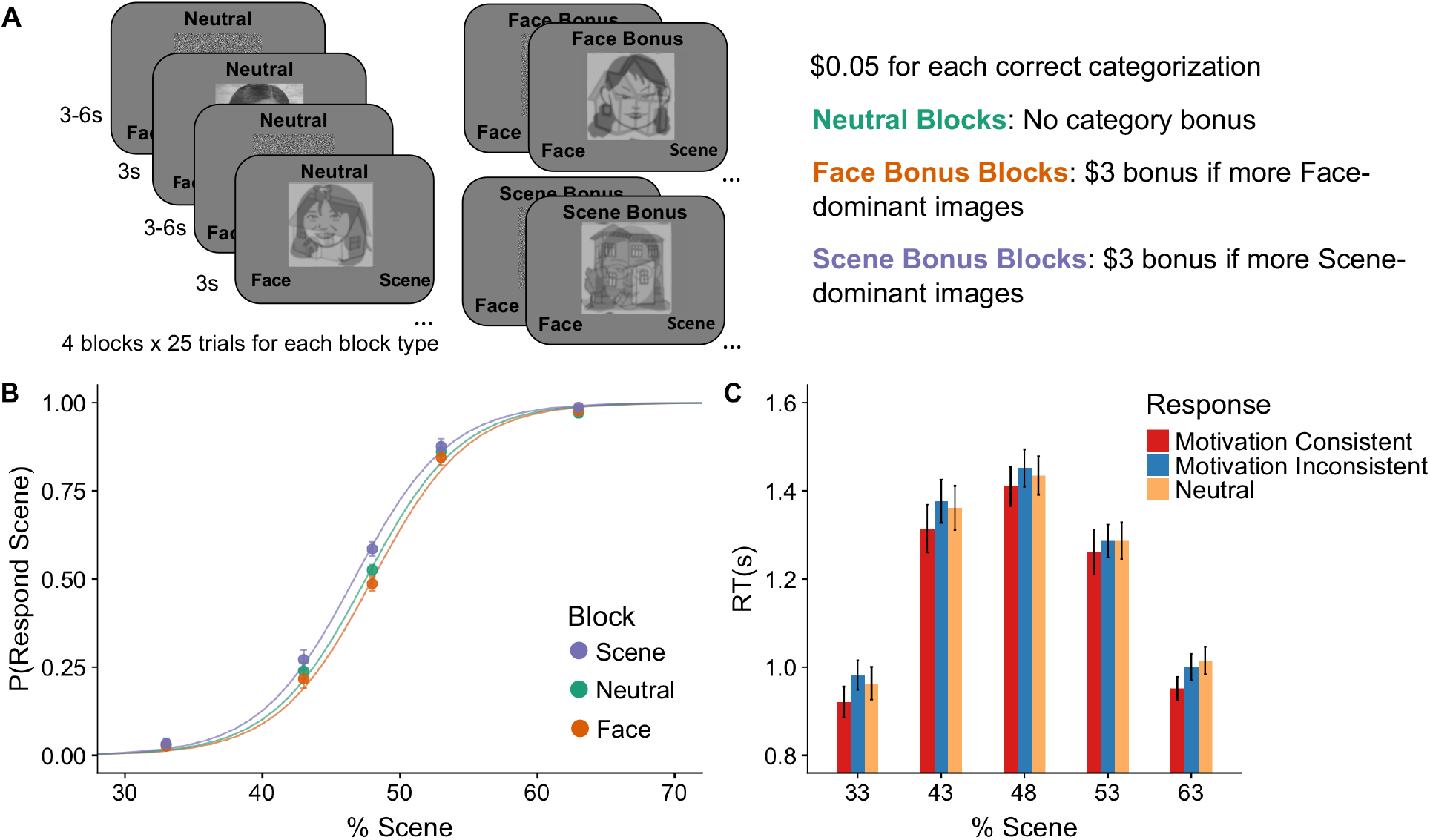
Motivation biased perceptual judgments. **A. Experimental Design.** Participants were rewarded for correctly categorizing ambiguous face/scene morphs as face-dominant or scene-dominant. On Face and Scene Bonus blocks, participants received a bonus if there were more face- or scene-dominant images in the block respectively. Images have been replaced with cartoon images to comply with *bioRxiv* policy. **B. Psychometric curves**. For the same level of objective evidence (% Scene) participants were more likely to categorize an image as scene-dominant on Scene Bonus Blocks and face-dominant on Face Bonus Blocks than on Neutral Blocks. **C. Response Times.** Responses that were motivation consistent (e.g., categorizing an image as face-dominant in a Face Bonus block) were faster than responses that were motivation inconsistent. Error bars indicate between-participant standard error of the mean.

Prior to the start of the experiment, participants were given explicit instructions that “the category bonus is based on the actual number of face-dominant or scene-dominant images in the block, and not on the categorizations that [they] make” and that “to earn more money, [they] should be as accurate as possible”. The instructions were delivered verbally by the experimenter, and also in written form on screen at the beginning of the experiment. As the instructions were intuitive and explicit, it is unlikely that participants misunderstood the task (see *Supplemental Material* for additional discussion). At the end of each Face Bonus or Scene Bonus block, participants received feedback on whether there were more face-dominant or scene-dominant images in the block and whether they earned the $3.00 bonus. For each participant, two Face-Bonus blocks contained one more face-dominant image, while the other two contained one less face-dominant image. Similarly, two Scene-Bonus blocks contained one more scene-dominant image while the other two contained one less scene-dominant image. Thus, each participant earned the $3.00 bonus on two Face-Bonus blocks and two Scene-Bonus blocks. Participants performed the blocks in a pseudo-randomized order such that they would not perform the same type of block consecutively.

### Behavioral Analyses

We modeled participants’ response data using generalized linear mixed-effects models (GLME), which allows for the modelling of all participants’ data in a single model rather than fitting a separate model for each participant (Knoblauch & Maloney, 2012). We modeled participants’ response (i.e. face-dominant or scene-dominant) as a function of the % scene in an image and block type (contrast coding with *Neutral* blocks as the reference condition). The model included random slopes and intercepts for % scene and block type to account for random effects across participants (M1; see Table S1 for full model specification). Model estimation was performed using the glmer function in the lme4 package in R (Bates et al., 2015), with *p*-values computed from t-tests with Satterthwaite approximation for the degrees of freedom, as implemented in the lmerTest package (Kuznetsova et al., 2019).

We ran two linear mixed effects models (LME) to examine the effect of motivation on participants’ response times (RT). RTs were modeled as a function of participants response type, that is whether participant’s response was *motivation consistent* (i.e. categorizing an image as the category they were motivated to see), *motivation inconsistent* (i.e. categorizing an image as the category they were motivated to not see) or *neutral* (i.e. trials in *Neutral* blocks). Both models coded response type using contrast coding but differed on which response type was coded as the reference group. The first model coded motivation consistent responses as the reference group, and tested whether RTs were significantly different for motivation consistent responses than for motivation inconsistent and neutral responses (M2; Table S1). The second model coded neutral responses as the reference group, and tested whether RTs were significantly different for neutral responses than for motivation consistent and motivation inconsistent responses (M3; Table S1). Both models controlled for whether participants categorized the image as face-dominant or scene-dominant. The models also included the absolute difference between % scene and % face in an image as a covariate of no interest to control for trial difficulty, as well as random intercepts and random slopes for all predictor variables.

### Pupillometry Analyses

Pupil diameter was recorded using an EyeLink 1000 system (SR Research) at a sampling rate of 500 Hz. The eye-tracker was calibrated using a standard 5-point calibration sequence. If calibration failed when recording data from both eyes, calibration was reattempted while recording from one eye (n = 12). If calibration remained unsuccessful, pupil data were not collected, and the participant was excluded from all analyses (n = 3). The eye-tracker was recalibrated half-way through the experiment. Eye-blinks and saccades were detected using Eyelink detection algorithms with default settings. Data were linearly interpolated from 150ms before the start of each blink or saccade to 150ms after the end of the blink or saccade. The resulting data were smoothed using a zero-phase low-pass filter (third-order Butterworth, cutoff = 4Hz) (Kret & Sjak-Shie, 2019), and z-scored for each eye and each participant. Data recorded from both eyes were then averaged to obtain a single time course.

We then extracted the timepoints of interest for each trial (stimulus-locked response: 500ms before stimulus onset to 2 seconds after stimulus onset; decision-locked response: 2 seconds prior to decision). Trials on which more than 30% of the data were missing were discarded (M = 1.8% of total trials, SE = 0.6%). For each participant, we modeled each timepoint using a general linear model:

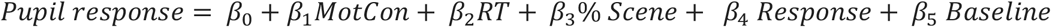

where *MotCon* was coded as 1 for motivation consistent responses (e.g., categorizing an image as scene-dominant in a Scene Bonus Block) and 0 for motivation inconsistent responses (e.g., categorizing an image as face-dominant in a Scene Bonus Block). *Response* was coded as 1 for scene-dominant responses and 0 for face-dominant responses. *Baseline* denotes the average pupil diameter in the 500ms before stimulus onset and was included to account for trial-by-trial differences in baseline pupil diameter when assessing the pupil response. The regression coefficients for *MotCon* thus reflect the extent to which the pupil response at a given timepoint was different when the participant made motivation consistent responses than when they made motivation inconsistent responses, while controlling for RT, percent scene in an image, whether participants categorized the image as face-dominant or scene dominant and baseline pupil diameter.

Significance testing and correction for comparisons over multiple timepoints were then performed using non-parametric cluster-based permutation tests (Maris & Oostenveld, 2007), separately for the stimulus-locked timepoints and the decision-locked timepoints. At each time-point, a one-sample t-test was performed to assess if the regression coefficient was different from zero across participants. Clusters were defined as contiguous timepoints where the *t*-test resulted in a *p*-value < 0.05. Cluster size was computed as the sum of *t*-values in the cluster. A null distribution of maximal cluster sizes was generated by repeating the cluster-forming procedure 10,000 times with data where the labels for motivation consistent and motivation inconsistent responses were randomly shuffled. Family-wise error rate corrected p-values were then determined as the proportion of the null distribution where the maximal cluster size was greater than the observed cluster size.

To examine the separate effects of baseline and evoked pupil dilation, we defined baseline pupil dilation as the average pupil diameter in the 500ms before stimulus onset, and the evoked pupil response as the average change in pupil diameter from baseline in the 1 second prior to the time of choice. We modeled whether participants made motivation consistent responses as a function of baseline pupil dilation, the evoked pupil response, RT and trial difficulty (i.e. absolute difference between % scene and % face of the image) using a linear mixed effects model (M4; Table S1). This analysis allowed us to assess the relationship between each pupil signal and participants’ responses while controlling for one another, RT and trial difficulty. We assessed the relationship between pupil dilation and trial accuracy and RT using a similar approach (M5 & M6; Table S1), and ran separate analyses to test for the zero-order relationship between baseline pupil diameter and motivation consistent responses (Supplemental Materials).

### Drift Diffusion Model

The drift diffusion model (DDM) is a class of sequential sampling models commonly applied to two-alternative forced choice paradigms (Ratcliff & McKoon, 2007). In the context of our task, a DDM assumes that participants’ responses arise from the noisy accumulation of sensory information (Fig. 2A). If the level of evidence crosses one of two decision thresholds (upper bound = scene; lower bound = face), the corresponding response is made. The starting point and rate of evidence accumulation were determined by the free parameters *z* and *v* respectively. The distance between the two thresholds was determined by the free parameter *a*, while time unrelated to the decision process (non-decision-time; e.g., time needed for motor response) was determined by the free parameter *t*.

**Figure 2.**
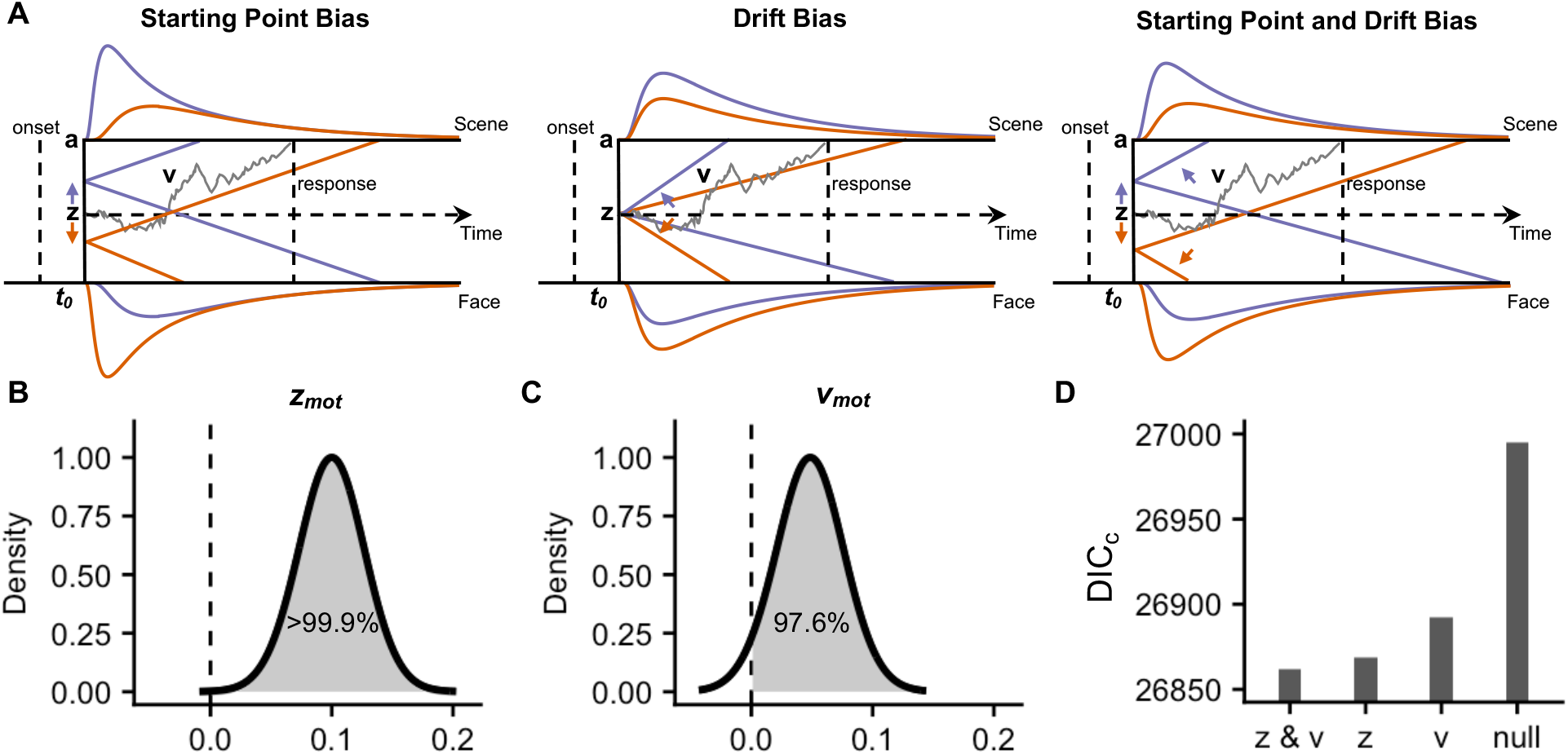
Motivation biased starting point and rate of evidence accumulation. **A. Schematic depiction of biasing mechanisms**. Biases in starting point and drift rate have distinguishable effects on the shape of response time (RT) distributions. Grey line: example trajectory of evidence accumulation on a single trial. Purple line: mean drift and resulting RT distribution when motivated to see scene-dominant images; Orange line: mean drift and resulting RT distribution when motivated to see face-dominant images. *z* = starting point, *v* = drift rate, *a* = threshold, *t_0_* = non-decision time. **B. Posterior distribution of model-estimated starting point bias (*z_mot_*)**. Dashed line indicates 0 (no bias). **C. Posterior distribution of model-estimated drift bias (*v_mot_*).** Dashed line indicates 0 (no bias). **D. Model comparison based on DIC_*c*_**. Model where motivation biased both the starting point and drift rate (*z & v*) yielded a lower DIC_c_ score than models where motivation biased only the starting point (*z*), only the drift rate (*v*) or neither parameters (*null*).

Model parameters were estimated from participants’ response time distributions using the HDDM toolbox with default priors (Wiecki et al., 2013; *Supplemental Material*). HDDM implements hierarchical Bayesian estimation, which assumes that parameters for individual participants were randomly drawn from a group-level distribution. We estimated group-level parameters as well as parameters for each individual participant, where each participant’s parameters both contributed to and were constrained by parameters at the group level. Markov chain Monte Carlo (MCMC) sampling methods were used to estimate the joint posterior distribution of all model parameters (30,000 samples; burn-in = 3,000 samples; thinning = 2). To account for outliers generated by a process other than that assumed by the model (e.g., lapses in attention, accidental button press), we estimated a mixture model where 5% of trials were assumed to be distributed according to a uniform distribution.

The HDDM package allows for parameters to vary according to a specified linear model. To examine the effects of motivation on the starting point, we allowed the starting point *z* to vary as a function of the motivation consistent category. HDDM implements *z* as the *relative* starting point, ranging from 0 to 1, with 0.5 denoting an unbiased starting point. As such, we used the inverse logit link function to restrict *z* to values between 0 and 1:

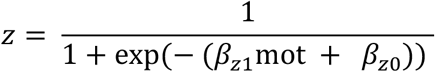

where *mot* denotes the motivation consistent category defined by the different types of experimental blocks. *mot* was coded as 1, −1, and 0 for Scene-Bonus blocks, Face-Bonus blocks and Neutral blocks respectively. Positive values of *β_z1_* reflect moving the starting point towards the Scene threshold on Scene-Bonus blocks, and towards the Face threshold on Face-Bonus blocks. We took *β*_*z*1_ as a measure of the motivational bias in the starting point (*z_mot_*).

In the same model, we modeled the drift rate *v* as a function of the motivation consistent category:

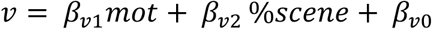

where *mot* was again coded as +1 (Scene-Bonus block), −1 (Face-Bonus Block) and 0 (Neutral Block). Positive values for *β*_*v*1_ reflect a drift bias towards the Scene threshold on Scene-Bonus blocks and towards the Face threshold on Face-Bonus blocks. We took *β*_*v*1_ as a measure of the motivational bias in drift rate (*v_mot_*). *β*_*v*2_ reflects the effect of sensory evidence (i.e. percentage scene of an image) on the drift rate. We demeaned % scene prior to entering it into the model such that the intercept term, *β*_*v*0_, would also reflect individual participant’s intrinsic biases in the drift rate at 48% scene (i.e. the point of subjective equivalence as established by pilot data).

For each of the bias parameters (*z_mot_* and *v_mot_*), we computed the proportion of the posterior distribution that was greater than 0. This proportion denotes the probability that the parameter had a positive value (i.e. a positive motivational effect on the parameter). To examine if either of the biases were sufficient for explaining the data, we fit two additional comparison models in which only *z* or only *v* was biased by motivation. As a baseline for comparison, we also fit a *null* model in which neither the starting point nor drift rate were biased by motivation. While HDDM models are commonly compared using deviance information criterion (DIC; Spiegelhalter et al., 2002; Wiecki et al., 2013), DIC is known to favor models with greater complexity (Plummer, 2008). We thus compared the four models using a corrected DIC measure (DIC_c_) that penalizes twice the number of effective parameters (Ando, 2011):

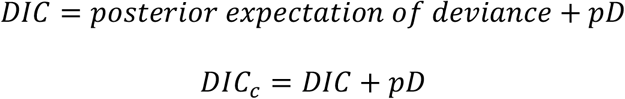

where *pD* denotes the number of effective parameters, with lower DIC_c_ values indicating better model fit. Model recovery simulations indicated that DIC_c_ accurately identifies the true model from simulated data (*Supplemental Material,* Fig. S2).

Next, we assessed how trial-by-trial fluctuations in pupil dilation relate to the two biasing mechanisms. We computed the evoked pupil response on each trial as the average change in pupil dilation from baseline in the 1 second prior to choice. This duration corresponded to the time period during which the response-locked pupil time course was higher on motivation consistent trials than on motivation inconsistent trials. As was the case in our earlier analyses, baseline pupil diameter was defined as the average pupil diameter in the 500ms prior to stimulus onset. To examine the relationship between the evoked pupil response and motivational bias, we generated a regressor that was the interaction between the evoked pupil response and the motivation consistent category:

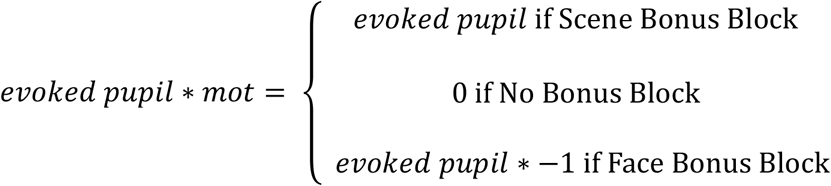

We generated a similar regressor, *baseline*mot*, to examine the relationship between baseline pupil diameter and motivational bias. We allowed the starting point to vary as a function of the *evoked pupil*mot* and *baseline*mot*:

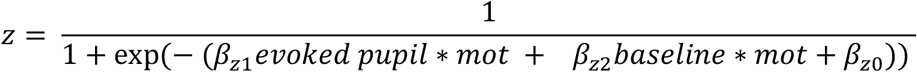

Positive values for *β_z1_* or *β_z2_* would indicate that the corresponding pupil signal has a positive relationship with the starting point on Scene Bonus blocks (i.e. reducing the amount of evidence needed to respond “Scene-dominant”) and a negative relationship on Face Bonus blocks (i.e. reducing the amount of evidence needed to respond “Face-dominant”). Even though the evoked pupil response occurred after the start of the trial, it can bias the estimate of the starting point. This is because a model with a biased starting point is functionally equivalent to a model with asymmetric decision thresholds. For example, a model in which the starting point is biased towards the Scene threshold is indistinguishable from a model with a lower Scene threshold and unchanged face threshold. As the HDDM implements symmetric thresholds with a relative starting point, the effect of asymmetric thresholds can only be captured as a biased starting point. The evoked pupil response can increase participants’ likelihood of making a motivation consistent choice by reducing the corresponding decision threshold (i.e. reducing the amount of evidenced needed to make that response). Including *evoked pupil*mot* in the model allowed us to test this possibility.

To examine the relationship between pupil dilation and motivational biases in the drift rate, we modeled the drift rate as a function of *evoked pupil*mot* and *baseline*mot*:

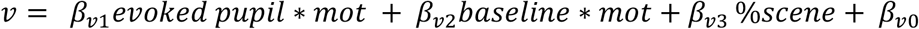

Positive values for *β_v1_* and *β_v2_* would indicate that corresponding pupil signal has a positive relationship with the drift rate (i.e. faster accumulation towards the scene threshold) on Scene Bonus blocks, and a negative relationship (i.e. faster accumulation towards the face threshold) on Face Bonus blocks.

Model convergence was formally assessed using the Gelman-Rubin 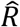 statistic (Gelman & Rubin, 1992), which runs multiple Markov chains to compare within-chain and between-chain variances. Large differences 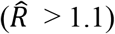 between these variances would signal non-convergence. In addition, we examined each trace to check that there were no drifts or large jumps, which would also suggest non-convergence. We report model convergence metrics, posterior means and 95% credible intervals of model parameters in Tables S2. We also performed posterior predictive checks by comparing participants’ behavior with data simulated with parameter values sampled from the posterior distribution estimated by each model (*Supplemental Material*).

## Results

Thirty-eight participants were presented with visually ambiguous images created by averaging a face image and a scene image in different proportions, and were rewarded for correctly categorizing whether the image was face-dominant or scene-dominant (Fig. 1A). Participants performed the task in blocks of 25 trials. In Face Bonus blocks, participants received a $3 category bonus if there were more face-dominant images in the block. In Scene Bonus blocks, participants received a $3 category bonus if there were more scene-dominant images in the block. In Neutral blocks, participants did not receive a category bonus. As such, participants were motivated to see more face-dominant images in Face Bonus blocks, and scene-dominant images in Scene Bonus blocks. Crucially, the category bonuses were determined by the objective composition of images in each block, and not by participants’ responses. To earn the most money, participants should ignore the category bonus and categorize the images accurately. Nevertheless, participants’ categorizations might be biased by what they were motivated to see.

### Motivation biased perceptual judgments

To assess the effects of the category bonuses on perceptual judgments, we estimated participants’ psychometric function separately for each block type (Fig. 1B). Statistical comparisons between the curves were performed using generalized linear mixed effects models (GLME). Relative to Neutral blocks, participants were more likely to categorize an image as face-dominant in Face Bonus blocks (*z* = 2.09, *p* = 0.037, b = 0.15, 95% CI = 0.01 to 0.28), and scene-dominant in Scene Bonus blocks (*z* = 3.21, *p* = 0.001, b = 0.25, 95% CI = 0.09 to 0.40). Thus, for the same face vs. scene proportion, participants were more likely to categorize the image as the category they were motivated to see.

Next, we assessed if motivation affected participants’ response times (Fig. 1C). Participants’ were faster to categorize an image as the category they were motivated to see (motivation consistent responses, e.g., categorizing an image as face-dominant in Face Bonus blocks) than as the category they were motivated not to see (motivation inconsistent responses; *t*(36.1) = 2.90, *p* = 0.006, *b* = 0.043, 95% CI = 0.006 to 0.054). Response times were faster for motivation consistent responses than on trials in the Neutral blocks ((*t*(36.1) = 2.48, *p* = 0.018, *b* = 0.031, 95% CI = 0.010 to 0.047), but were not significantly different between motivation inconsistent responses and trials in the in the Neutral blocks (*t*(34.2) = 1.01, *p* = 0.321, *b* = 0.013, 95% CI = −0.012 to 0.038).

### Motivation biases both starting point and rate of sensory evidence accumulation

We fit a drift diffusion model (DDM) to participants’ response time distributions. The DDM assumes that each perceptual judgment arises from the noisy accumulation of sensory evidence towards one of two decision thresholds, and provides a computational description of this process. Within the DDM framework, participants’ motivational bias can reflect a bias in the starting point (*z*) and/or a bias in the rate (*“drift rate”*, *v*) of sensory evidence accumulation in favor of the category participants were motivated to see. A bias in the starting point reduces the amount of evidence needed to make a motivation consistent response while a bias in the drift rate biases the evidence accumulation process in favor of motivation consistent category. Both biases increase the proportion of motivation consistent responses, but have distinguishable effects on the distribution of response times (Leong et al., 2019; White & Poldrack, 2014; *Supplemental Material*; Fig. 2A, Fig. S3). We estimated the extent to which each bias contributed to participants’ behavior by fitting the model to participants’ responses and response times (see *Methods*).

We allowed the starting point to vary according to a linear regression model with the motivation consistent category as a predictor variable. The regression coefficient reflects the extent to which motivation affects the starting point. Motivation had a positive effect on the starting point (*p*(*z_mot_* > 0) > 0.999, mean = 0.100, 95% credible interval = 0.053 to 0.145; Fig. 2B), indicating that the starting point was biased towards the scene threshold when participants were motivated to see scene-dominant images, and biased towards the face threshold when participants were motivated to see face-dominant images.

The drift rate was similarly modeled using linear regression. As the drift rate also depends on the amount of sensory evidence available in the image, we included the percentage scene in an image as an additional predictor. Motivation had a positive effect on the drift rate (*p*(*v_mot_* > 0) = 0.976, mean = 0.048, 95% credible interval = 0.0004 to 0.094, Fig. 2C), indicating that sensory evidence accumulated more quickly for the motivation consistent category. As expected, percentage scene had a positive effect on the drift rate (*p*(*v_scene_* > 0) > 0.999, mean = 1.446, 95% credible interval = 1.325 to 1.569), indicating that sensory evidence accumulation was biased towards the scene threshold at high scene proportion and biased towards the face threshold at low scene proportion.

To examine if either biasing mechanism was sufficient to account for participants’ data, we fit additional models where motivation biased only the starting point (*z* model) or only the drift rate (*v* model). Model fits were then compared using DIC_c_, with lower values indicating better fit (see *Methods*). The model in which motivation biased both the starting point and drift rate yielded the lowest DIC_c_ value (DIC_c_ - *z* & *v*: 26862, *z*: 26869, *v*: 26892, null: 26995; Fig. 2D). Furthermore, simulated data generated by parameterizing the *z & v* model with best-fit parameter values aligned well with participants’ data, and matched qualitative patterns in the data better than the alternative models (*Supplemental Material,* Fig. S3). Taken together, our modeling results suggest that motivation biased both the starting point and rate of sensory evidence accumulation in favor of the motivation consistent category.

### Pupil-linked arousal was higher when making motivation consistent responses

We next investigated whether physiological arousal was associated with motivational biases in perceptual judgments. We measured participants’ pupil diameter as a measure of physiological arousal. For each trial, the pupil time course was extracted around two events of interest i. stimulus onset (500ms before stimulus onset to 2 seconds after stimulus onset; Fig. 3A) and ii. choice (2 seconds prior to choice; Fig. 3B). We then averaged the pupil time courses separately for trials on which participants made motivation consistent responses and trials on which they made motivation inconsistent responses. The shapes of the pupil time courses were similar across the two types of trials. In particular, stimulus onset induces an increase in pupil diameter that peaks around 300 ms, reflecting an increase in arousal at the start of the trial. This is followed by a brief recovery before a second increase that peaks around 250ms post-choice.

**Figure 3.**
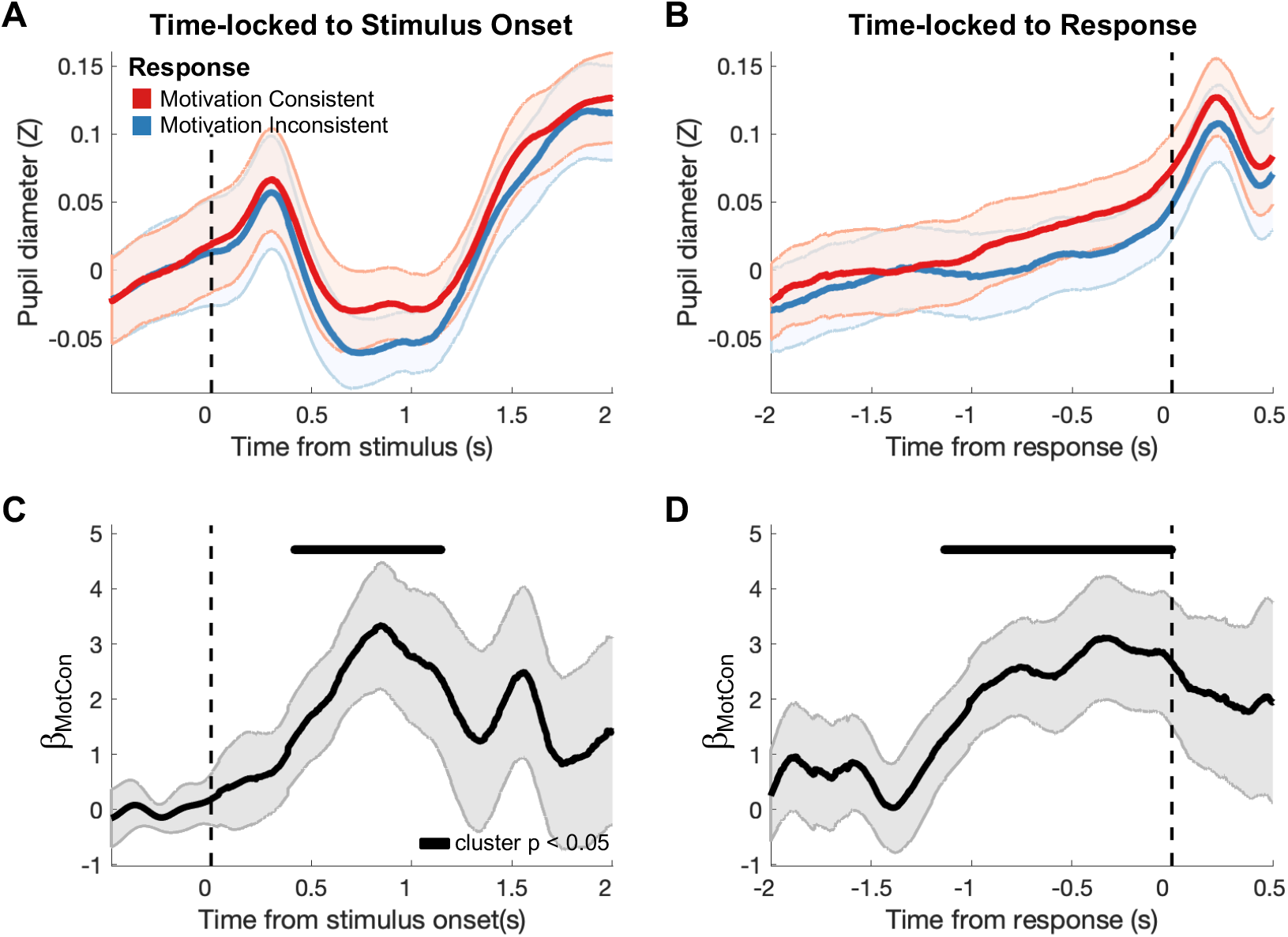
Motivation consistent responses are associated with greater pupil dilation. **(Top)** Average pupil diameter over the course of a trial, time-locked to **A.** stimulus onset and **B.** time of response. Solid lines denote average pupil time course for trials on which participants made motivation consistent responses (red) and motivation inconsistent responses (blue). Shaded error bars denote between-participant S.E.M. (**Bottom)** Regression coefficient indicating the relationship between making a motivation consistent response and pupil dilation, controlling for RT, percent scene, whether participants made a face-dominant or scene-dominant categorization, and baseline pupil diameter, time-locked to **C.** stimulus onset and **B.** time of response. Shaded error bars denote S.E. of the estimate. Black horizontal bars indicate time points where regression coefficients were significantly different from 0 based on cluster-based permutation tests.

We modeled the pupil time courses using a general linear model (GLM) approach (see *Methods*), which allowed us to examine the relationship between motivation consistent responses and pupil dilation, while controlling for the effects for RT, percent scene in an image, whether participants categorized the image as face-dominant or scene-dominant, and baseline pupil diameter. Using this approach, we found that pupil size was larger on trials where participants made motivation consistent responses than on trials where they made motivation inconsistent responses (from 400ms to 1.1 seconds after stimulus onset, cluster *p* = 0.028; from 1.1s prior to decision to time of decision, cluster *p* = 0.008; Fig. 3C, Fig. 3D). These results indicate that pupil-linked arousal was associated with motivational biases in perceptual judgments.

To examine the role of both baseline and evoked pupil dilation, we defined baseline pupil dilation as the average pupil diameter in the 500ms before stimulus onset, and the evoked pupil response as the average change in pupil diameter from baseline in the 1 second prior to the time of choice. We then entered both measures into linear mixed effects models to assess their relationship with motivation consistent responses, accuracy and RT (see *Methods*). Reproducing the time course analyses, participants were more likely to make motivation consistent responses on trials with higher evoked pupil response (*z* = 2.89, *p* = 0.004, *b* = 0.166, 95% CI = 0.054 to 0.278), controlling for the effects of RT, baseline pupil diameter and trial difficulty. Participants were also less likely to correctly categorize the image as face-dominant or scene-dominant on trials with higher evoked pupil response (*z* = −2.46, *p* = 0.014, *b* = −0.388, 95% CI = −0.698 to −0.080). Together, these results suggest that at higher levels of arousal, participants’ perceptual judgments were more likely to be biased by the motivation manipulation and depended less on the objective information in the image.

Consistent with earlier work (de Gee et al., 2014), evoked pupil responses were larger on trials with longer RTs (*t*(36.8) = 2.84, *p* = 0.007, *b* = 0.137, 95% CI = 0.043 to 0.231). As reported earlier, motivation consistent responses were on average faster than motivation inconsistent responses. If the relationship between pupil dilation and motivation consistent responses were driven solely by RT effects, the shorter RTs on motivation consistent responses should have led to a smaller evoked pupil response, which is the opposite of what was observed. As such, RT effects cannot explain the relationship between the evoked pupil response and motivational biases in perceptual judgments.

In contrast, baseline pupil diameter was not associated with motivation consistent responses (*z* = 0.998, *p* = 0.318, *b* = 0.028, 95% CI = −0.026 to 0.082), accuracy (*z* = −0.665, *p* = 0.506, *b =* −0.040, 95% CI = −0.158 to 0.078) and RTs (*t(31.5)* = 1.40, *p* = 0.172, *b =* 0.018, 95% CI = −0.007 to 0.042). The preceding analyses assessed the effects of baseline pupil diameter while controlling for the evoked pupil response and trial difficulty. We ran a separate analysis to test for the zero-order relationship between baseline pupil diameter and motivation consistent responses, which similarly found that baseline pupil diameter was not different between motivation consistent and motivation inconsistent responses (*t(37.3)* = −0.009, *p* = 0.993, *b =* 0.0002, 95% CI = −0.049 to 0.049; Fig S4). These results indicate that the relationship between pupil dilation and motivation consistent responses was driven by larger evoked pupil responses during the decision period and not trial-by-trial fluctuations in the pre-stimulus baseline.

In exploratory analyses, we found that baseline pupil diameter was marginally lower for neutral responses (i.e. trials in the *Neutral* blocks without a $3 category bonus) than both motivation consistent and motivation inconsistent responses (neutral vs. motivation consistent: *t(37.1)* = 1.76, *p* = 0.088, *b =* 0.072, 95% CI = −0.001 to 0.152; neutral vs. motivation inconsistent: *t(37.1)* = 1.86, *p* = 0.071, *b =* 0.072, 95% CI = −0.004 to 0.148; Fig. S4). This suggests that the prospect of a category bonus induced an increase in baseline pupil diameter, though the raised baseline on its own did not bias participants to make motivation consistent responses. Instead, they suggest that the bias was associated with a larger evoked response on top of the elevated baseline.

In summary, increased pupil dilation during the decision period was associated with participants making motivation consistent perceptual categorizations. This relationship was not due to differences in RT or baseline pupil diameter. In addition, increased pupil dilation during the decision period was associated with lower task accuracy, suggesting that on trials with heightened arousal, participants were more biased by what they were motivated to see and relied less on the objective information in the image.

### Pupil-linked arousal was associated with trial-by-trial motivational biases in drift rate

Pupil-linked arousal was associated with an increased likelihood of making motivation consistent perceptual judgments. Our earlier modeling analyses suggest that motivational effects on perceptual judgments were driven by both biases in the starting point and rate of evidence accumulation. Was pupil-linked arousal related to either or both biasing mechanisms? To address this question, we used a linear regression approach to examine the relationship between pupil dilation and trial-by-trial fluctuations in motivational bias. Instead of estimating a fixed starting point or drift rate across trials, we allowed the two parameters to vary according to the baseline pupil diameter and the evoked pupil response on each trial.

To model the relationship between pupil dilation on motivational biases, we generated regressors that denote the interaction between pupil dilation and the motivation consistent category (i.e. *baseline*mot* and *evoked pupil*mot*; see Methods). The coefficients on these regressors reflect the extent to which the corresponding pupil signal (i.e. baseline diameter or evoked pupil response) was associated with motivational effects on model parameters. For example, a positive regression coefficient for *evoked pupil*mot* on the drift rate would reflect a positive relationship between the evoked pupil response and motivational biases in the drift rate. We fit participants’ data to a model where the starting point and drift rate were allowed to vary as a function of *baseline*mot* and *evoked pupil*mot*.

The evoked pupil response had a positive relationship with motivational biases in the drift rate (*p*(*v_evoked*mot_* > 0) = 0.979, mean = 0.069, 95% credible interval = 0.004 to 0.135; Fig. 4A). That is to say, larger evoked pupil responses were associated with a bias in evidence accumulation towards the scene threshold when participants were motivated to see scene-dominant images and a bias towards the face threshold when they were motivated to see face-dominant images. In contrast, the evoked pupil response was not associated with motivational biases in the starting point (*p*(*z_evoked*mot_* > 0) = 0.437, mean = −0.003, 95% credible interval = −0.082 to 0.089). Baseline pupil diameter was not associated with motivational biases in either the drift rate (*p*(*v_baseline*mot_* > 0) = 0.851, mean = 0.018, 95% credible interval = −0.015 to 0.058) or the starting point (*p*(*z_baseline*mot_* > 0) = 0.366, mean = −0.006, 95% credible interval = −0.043 to 0.031). Similar results were obtained with a model that included the motivation consistent category as a covariate (Fig. S5), and a model that included parameters for inter-trial variability of the starting point, drift rate and non-decision time (Fig. S6).

**Figure 4.**
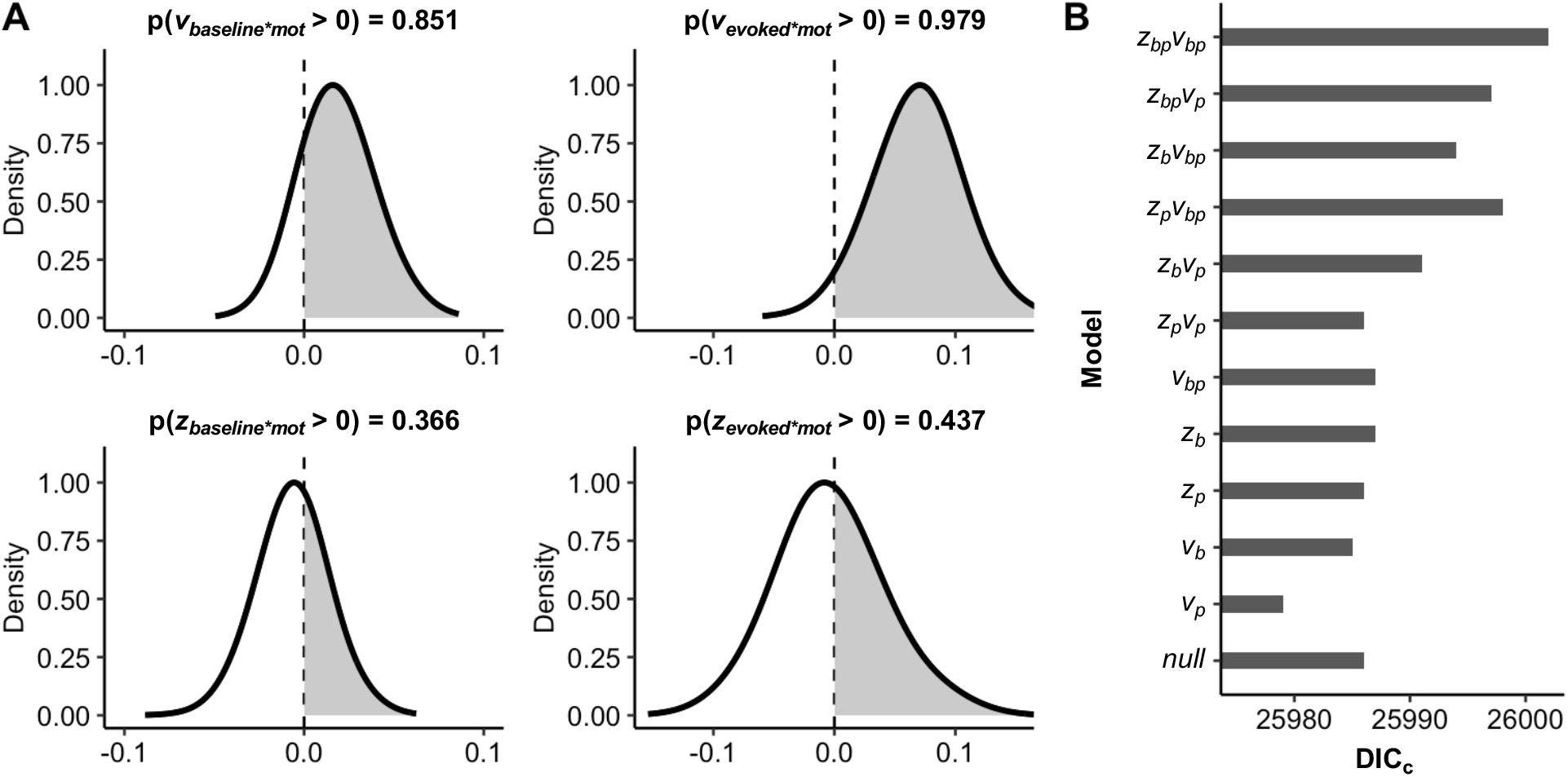
Motivational biases in drift rate are associated with trial-by-trial fluctuations in evoked pupil response. **A. Posterior distributions of regression coefficients.** Posterior distributions for the regression coefficients of baseline pupil diameter on drift bias (*v_baseline*mot_*) and starting point bias (*z_baseline*mot_*), and the regression coefficients of the evoked pupil response on drift bias (*v_evoked*mot_*) and starting point bias (z_*evoked*mot*_). Dashed line indicates 0 (no effect). There was a positive relationship between evoked pupil response and motivational biases in the drift rate (*p*(*v_evoked*mot_* > 0) > 0.95). **B. Model comparison based on DIC_c_.** *z* and *v* denote models where pupil dilation modulated starting point and drift bias respectively. Subscript *b* indicates that the parameter varied with baseline pupil diameter while subscript *p* indicates that the parameter varied with evoked pupil response. *null* denotes a model where neither the starting point nor the drift rate varied with pupil dilation. A model where motivational biases in the drift rate varied with evoked pupil response (model *v_p_*) provided the best fit to data.

Our results suggest that the relationship between pupil dilation and motivational biases arose due to a bias in drift rate associated with larger evoked pupil responses during the decision period. If this were true, a simpler model where only the motivational bias in the drift rate varied as a function of *evoked pupil*mot* would provide a better fit to the data. Indeed, such a model had a lower DIC_c_ value than more complex models, models where other parameters were allowed to vary with pupil dilation, and a model where neither the starting point nor the drift rate were modulated by pupil dilation (Fig. 4B; *Supplemental Material*, Table S3). Furthermore, data simulated from a model where only the motivational bias in the drift rate varied as a function of *evoked pupil*mot* was sufficient to reproduce the empirical observation that evoked pupil responses were higher for motivation consistent responses than motivation inconsistent responses (empirical data: mean difference = 0.023, 95% CI = 0.002 to 0.444, *t*(37) = 2.24, *p* = 0.031; model *v_p_*: mean difference = 0.020, 95% CI = 0.009 to 0.300, *t*(37) = 3.87, *p* < 0.001; *Supplemental Material*, Fig. S7).

Together, the estimated regression coefficients from the model and model comparison using DIC_c_ provide converging evidence that arousal effects on motivational biases in perceptual judgments were driven by biases in evidence accumulation that tracked trial-by-trial fluctuations in evoked pupil response.

## Discussion

In the current work, we investigated the role of physiological arousal in modulating motivational biases in perceptual decision-making. We manipulated the percept participants were motivated to see as they performed a perceptual decision-making task. Participants were more likely to categorize an image as the category they were motivated to see. This motivational bias reflected both an *a priori* bias towards the motivation consistent category, as well as a bias in how participants accumulated sensory evidence over time. Arousal, as measured by pupil dilation, was higher when participants made motivation consistent responses. Trial-by-trial fluctuations in the evoked pupil response were specifically associated with faster accumulation of sensory evidence in favor of the motivationally desirable percept. These findings suggest that pupil-linked arousal processes mediate motivational effects on perceptual decision-making by enhancing the processing of motivationally desirable information.

Contemporary accounts of perceptual decision-making propose that perceptual decisions are determined by comparing the activity of neurons selective to different perceptual features (Gold & Shadlen, 2007; Heekeren et al., 2008). Motivation can bias this comparison in favor of percepts participants are motivated to see by enhancing neural activity selective to desirable perceptual features in sensory regions of the brain (Leong et al., 2019). For example, when presented with ambiguous face-scene image morphs, face-selective and scene-selective neural activity in sensory cortices was greater when participants were motivated to see face-dominant and scene-dominant images respectively. Participants with stronger motivational enhancement of category-selective neural activity also exhibited greater biases in model-estimated drift rate, suggesting that motivationally enhanced neural representations result from the biased accumulation of sensory information. The current findings suggest motivational biases in sensory processing not only vary between individuals, but also vary trial-by-trial depending on the level of arousal. When arousal is high, there is a stronger motivational bias in sensory processing and participants are more likely to see the desirable percept. In contrast, when arousal is low, motivational effects are weaker and participants’ decisions depend more on the objective sensory information in the image.

Pupil-linked arousal processes are thought to be driven by activity in the locus coeruleus norepinephrine system (Joshi et al., 2016; Murphy et al., 2014). Our results are consistent with the GANE model of norepinephrine function, which posits that arousal-related norepinephrine release selectively enhances the processing of salient stimuli (Mather et al., 2016). The GANE model builds on earlier work showing that norepinephrine increases the *gain* of neurons, such that excited neurons become even more active and inhibited neurons become even less active (Aston-Jones & Cohen, 2005; Sara & Bouret, 2012). High neural gain would thus result in focused attention on the most salient features of a stimulus (Eldar et al., 2016). Here, we argue that otherwise neutral perceptual features become *motivationally salient* when participants are motivated to see them. Attention is directed towards these features, and neurons encoding these features would be more active than neurons encoding other features (Bourgeois et al., 2016; Pessoa, 2009). During moments of heightened arousal, high neural gain accentuates this difference, selectively enhancing the neural representation of motivationally salient features and amplifying motivational effects on perceptual processing. Novel imaging approaches now provide the opportunity to accurately and reliably measure activity in the locus coeruleus (Betts et al., 2019; Liu et al., 2017). Future work can take advantage of these approaches to simultaneously measure activity in the locus coeruleus and sensory regions of the brain to directly test the role of the locus coeruleus norepinephrine system in enhancing motivationally desirable neural representations.

Growing evidence suggests that pupil-linked arousal processes during the decision period (i.e. between stimulus onset and making a response) influences the accumulation of sensory evidence during perceptual decision-making (Cheadle et al., 2014; de Gee et al., 2014, 2017, 2020; Keung et al., 2019; Krishnamurthy et al., 2017). For example, in the studies by de Gee and colleagues (2014; 2017; 2020), larger evoked pupil responses were associated with a reduction in participants’ intrinsic biases to report the presence or absence of a target pattern embedded in a noisy sensory background. Similarly, Krishnamurthy and colleagues (2017) found that larger evoked pupil responses were associated with the reduced influence of prior expectations in a sound localization task. How might we reconcile the negative relationship between pupil dilation and decision biases documented in these earlier studies with our finding that larger pupil dilation was associated with stronger motivational biases? One possible explanation is that in the earlier studies, participants were motivated to be accurate in their perceptual reports and were indifferent to the different perceptual options. As such, attention would not be biased towards any one aspect of the stimulus. The higher neural gain associated with a larger evoked pupil response would then primarily serve to increase neuronal sensitivity to the stimulus. Thus, larger evoked pupil responses would be associated with perceptual decisions that depend more on sensory features of the stimulus and less on prior biases. In contrast, participants in our task were motivated to see one of two competing percepts present in an ambiguous stimulus. Attention directed towards stimulus features associated with the desirable percept enhances neural activity encoding that percept. Increasing neural gain in our task amplifies the motivational enhancement and biases perceptual decisions in favor of the percept participants were motivated to see. Thus, larger evoked pupil responses would be associated with stronger motivational biases.

Prior studies have found that pre-stimulus arousal levels, as measured by pre-stimulus baseline pupil diameter, can also affect the upcoming decision (Krishnamurthy et al., 2017; Murphy et al., 2014; Nassar et al., 2012). For example, Krishnamurthy and colleagues (2017) found that both baseline and evoked pupil dilation were associated with the reduced influence of prior expectations on perceptual decisions. In our task, baseline pupil diameter was not related to motivational biases. Our task, however, was not designed to examine pupil dilation during the pre-stimulus period. In particular, we have a relatively short inter-trial-interval between the end of one trial and stimulus onset on the next trial (mean ITI = 3.5 seconds). As such, the pre-stimulus baseline on a given trial might be contaminated by the evoked pupil response on the previous trial. Future experiments with longer intervals between trials will be necessary to determine the relationship (or lack of) between baseline pupil diameter and motivational bias.

Altogether, the current results expand our understanding of the role of arousal in perceptual decision-making. While prior work has primarily focused on how arousal affects perceptual decision-making in “neutral” contexts where participants are indifferent to different perceptual options, our study examined the effects of arousal-related biases when participants are motivated to see one percept over another. Our findings suggest that arousal biases evidence accumulation in favor of desirable percepts. As a result, during moments of heightened arousal, participants were more likely to be biased to see what they were motivated to see and less likely to make accurate perceptual categorizations. Given the putative relationship between pupil-linked arousal processes and the locus coeruleus norepinephrine system, our results also highlight potential neuromodulatory mechanisms driving motivational biases. Notably, motivation has also been shown to influence information processing across many domains of human cognition (Kunda, 1990). For example, people learn more from positive outcomes than negative outcomes during an instrumental learning task (Lefebvre et al., 2017), and incorporate favorable information more than unfavorable information when updating beliefs about future life events (Sharot et al., 2011). Do pupil-linked arousal processes also modulate motivational biases beyond sensory perception? Given the wide-spread projections of norepinephrine neurons across the brain, this is certainly possible. Future studies can extend our work and examine the relationship between arousal and motivational biases on other types of human reasoning and evaluation processes.

## Open Practices Statement

The study reported in this article was not formally preregistered. De-identified data and analysis scripts can be accessed at [to be filled in on publication]. Requests for other experimental materials can be sent via email to the lead author.

