## Supplemental Material for "Pupil-linked arousal biases evidence accumulation towards desirable percepts during perceptual decision-making"

### Determining the point of subjective equivalence

To determine the point of subjective equivalence of the composite face-scene stimuli, we collected pilot data from 30 participants performing 40 trials of a face-scene classification task with similar composite face/scene stimuli across a range of % scene (1×100% scene, 3×65% scene, 5×60% scene, 7×55% scene, 8×50% scene, 7×45% scene, 5×40% scene, 3×35% scene and 1×0% scene). Participants were not motivated to see one category over another, thus these data allowed us to examine how participants classified these images in the absence of motivational biases.

We modeled participants' behavioral data using a generalized linear mixed-effects model (GLME) and estimated the point of subject equivalence (PSE) as the % scene at which participants were equally likely to categorize a stimulus as face-dominant or scene-dominant (Fig. S1). The model-estimated PSE was 48% scene. Hence, for the current experiment, we considered images with greater than 48% scene as scene-dominant, and images with less than 48% scene as face-dominant.

### Assessing participants' understanding of task instructions

Participants were given explicit instructions that "the category bonus is based on the actual number of face-dominant or scene-dominant images in the block, and not on the categorizations that [they] make" and that "to earn more money, [they] should be as accurate as possible". The instructions were delivered verbally by the experimenter, and also in written form on screen prior to the start of the experiment. As the instructions were intuitive and explicit, it is unlikely that participants misunderstood the task.

Participants were also given feedback on whether they earned the $3.00 bonus at the end of each block. If participants had misunderstood the instructions and thought that the $3.00 category bonus depended on their choices, the rational strategy would be to report at least more than half of the images as the category consistent with the bonus. Participants would quickly encounter blocks where they made more bonus-consistent categorizations but did not earn a category bonus, and thus realize their mistake. When analyses were restricted to trials from the second half of the task, the proportion of scene-dominant responses remained higher on Scene Bonus blocks than Face Bonus blocks ($b = 0.343$, SE = 0.144, z = 2.39, $p = 0.017$), suggesting that the motivational bias observed in our task was not driven by participants who misunderstood the instructions early in the experiment.

Participants completed a post-experiment questionnaire which contained an open-ended question asking if they had understood the instructions correctly, to which all participants responded “Yes” without additional qualifiers. While it is possible that participants incorrectly assumed that they had understood the instructions, this suggests that none of the participants felt confused by the task. Participants were also asked if they were influenced by the category bonuses. We recoded participants’ open-ended responses to “No”, “A little”, “Sometimes” and “Yes” (see Table S4 for raw responses). Out of 38 responses, 13 were coded as “No”, 5 were coded as “A little”, 4 were coded as “Sometimes” and 16 were coded as “Yes”. 3 of the 20 responses that were coded as “Sometimes” or “Yes” were qualified with statements indicating that they understood the task instructions (e.g., “Yes, so I tried to not read the name of the categories in order to not be biased”). 2 of 18 responses coded as “No” or “A little” included similar qualifiers (e.g., Not really, since my answers do not change whether or not I got paid.”). Interestingly, the proportion of scene-dominant responses was higher on Scene Bonus blocks than Face Bonus blocks even when analyses were restricted to the 18 participants who answered “No” or “A Little” ($b = 0.266$, SE = 0.106, z = 2.52, $p = 0.012$). This result argues against the interpretation that our findings were driven by a conscious strategy to report more motivation consistent categorizations due to misunderstanding the task instructions.

We report the psychometric functions of individual participants in Figure S8. Figure S9 shows the regression coefficients of block type on the proportion of scene-dominant responses when a separate GLM was fit to each participant’s data. 30 out of 38 participants had a positive regression coefficient (i.e. higher proportion of scene-dominant responses on Scene Bonus blocks than Face Bonus blocks). At the group level, the motivational bias remained significant when the three most extreme participants (participants 4, 9 and 28) were excluded from the analyses ($b = 0.278$, SE = 0.066, $z = 4.22$, $p < 0.001$), indicating that our results were not driven by extreme participants. Taken together, these data and analyses suggest that it is highly unlikely that our results were driven by some or all participants misunderstanding the task instructions.

**Model priors**

Model parameters for the drift diffusion model were estimated from participants' response time distributions using the HDDM toolbox with default priors (Wiecki et al., 2013). For a given participant $i$, parameters are assumed to be drawn from the following distributions:

$$a_i \sim Gamma(\mu_a, \sigma_a^2)$$

$$t_i \sim N(\mu_t, \sigma_t^2)$$

$$z_i = invlogit\big(N(\mu_t, \sigma_z^2\ )\big)$$

$$v_i = N(\mu_v, \sigma_v^2)$$

with hyper-priors:

$$\mu_a \sim Gamma(1.5, 0.75)$$

$$\mu_t \sim Gamma(0.4, 0.2)$$

$$\mu_z \sim N(0.5, 0.5)$$

$$\mu_v \sim N(2, 3)$$

$$\sigma_a \sim HN(0.1)$$

$$\sigma_t \sim HN(1)$$

$$\sigma_z \sim HN(0.05)$$

$$\sigma_v \sim HN(2)$$

where $Gamma$ denotes a gamma distribution parameterized by mean and standard deviation, $N$ denotes a normal distribution parameterized by mean and standard deviation, $invlogit$ denotes an inverse logit transform and $HN$ denotes a half-normal parameterized standard deviation.

For the regression-based analyses, we used the prior for the original parameter for the intercept, and non-informative priors for the regression coefficients.

**Model recovery simulations**

We ran model recovery studies to examine if $DIC_c$ accurately recovers the true model from data simulated from i. a model where motivation does not bias participants' perceptual judgments (*null* model), ii. a model where motivation biases participants' starting point (*z* model), iii. a model where motivation biases participants' drift rate (*v* model) and iv. a model where motivation biases both participants' starting point and drift rate ($z$ & $v$ model). For each model, we generated 25 datasets, each consisting of 38 simulated participants performing 100 trials while motivated to see face-dominant images (Face Bonus block), 100 trials while motivated to see scene-dominant images (Scene Bonus block) and 100 trials with no motivation manipulation (Neutral block). For the *null* model, participants' parameters were randomly drawn from the following distributions:

$$a \sim Gamma(2, 0.3)$$

$$t \sim N(0.5, 0.15)$$

$$z = invlogit\big(N(0.03, 0.04)\big)$$

$$v = N(0.06, 0.2)$$

These prior distributions are created to match parameter values estimated from fitting the *null* model to behavioral data.

For the *z* model, we implemented a bias in starting point as follows:

$$z_{int} = N(0.03, 0.04)$$

$$z_{mot} = N(0.15, 0.1)$$

$$z = \begin{cases} invlogit(z_{int} + z_{mot}) \text{ if Scene Bonus Block} \\ invlogit(z_{int}) \quad \text{if Neutral Block} \\ invlogit(z_{int} - z_{mot}) \text{ if Face Bonus Block} \end{cases}$$

where $z_{int}$ determines the starting point on Neutral blocks and $z_{mot}$ determines a starting point bias that would give rise to a ratio of motivation consistent and motivation inconsistent responses comparable to that observed in the behavioral data.

For the *v* model, we implemented a bias in drift rate as follows:

$$v_{int} = N(0.06, 0.2)$$

$$v_{mot} = N(0.1, 0.1)$$

$$v = \begin{cases} v_{int} + v_{mot} \text{ if Scene Bonus Block} \\ v_{int} \quad \text{if Neutral Block} \\ v_{int} - v_{mot} \text{ if Face Bonus Block} \end{cases}$$

where $v_{int}$ determines the drift rate on Neutral blocks and $v_{bias}$ determines a drift bias that would give rise to a ratio of motivation consistent and motivation inconsistent responses comparable to that observed in the behavioral data.

For the *z & v* model, we implemented both starting point and drift biases, but reduced the magnitude of each bias by approximately half such that the two biases together give rise to a ratio of motivation consistent and motivation inconsistent responses comparable to that observed in the behavioral data:

$$z_{mot} = N(0.1, 0.1)$$

$$v_{mot} = N(0.05, 0.1)$$

We then fit all four models (5000 samples; burn-in = 500 samples; thinning = 2) to each dataset and compared model fits using $DIC_c$ (Ando, 2011, see Methods). Across the datasets generated by each model, $DIC_c$ correctly identified the true model in 93.0% (SE = 3.4%) on average (Fig. S2). These results indicate that the four models are identifiable, and support the use of $DIC_c$ as a metric for model selection.

**Posterior predictive checks**

For each model, we simulated 100 datasets with parameter values sampled from the posterior distribution estimated by the model. Each simulated dataset comprised the same number of participants performing the same number of trials as the real dataset, and reflect the pattern of choice and response time data if the behavior of participants' in our task were perfectly described by the model. We overlay the response time distribution generated by a model where motivation biased both the starting point and drift rate with the true response time distributions, separately for face and scene responses and for each block to assess model fit (Fig. S3A).

A motivationally biased starting point and a motivationally biased drift rate both contribute to a greater proportion of motivationally consistent responses, but have distinguishable effects on the shape of the response time distributions. In particular, the effect of a biased starting point is strongest early on in a trial, and diminishes with the accumulation of sensory evidence. As such, a biased starting point has a disproportionate effect on fast responses. In contrast, a biased drift rate affects evidence accumulation at each time-step, and thus has a strong effect on both fast and slow responses. This is consistent with the interpretation that a biased starting point reflects an *a priori* bias to make motivation consistent responses that can be overcome as participants accumulate more information about the stimulus, while a biased drift rate reflects a bias in evidence accumulation that affects how new information is processed.

The distinct effects of the two biases can be visually examined by comparing the response proportions across conditions at different response time quantiles (conditional response time functions, CRT; White & Poldrack, 2014). We binned the response times for each participant into quartiles, and computed the proportion of trials on which participants responded scene in each quartile. The proportions were then averaged across participants separately for Face Bonus, Neutral and Scene Bonus blocks. The analysis was restricted to trials at 48% scene, as there were insufficient trials at the other levels of percent scene to be divided into quartiles for each experimental block type. We compared the CRT of simulated data generated by models where motivation biased either or both the starting point and drift rate to examine which model best reproduced the CRT computed from participants' data (Fig. S3B). Each model was parameterized with the best-fit parameters obtained when the corresponding model was fitted to participants' data

For the model where motivation biased only the starting point, the predicted proportion of scene responses in Scene Bonus blocks was higher than that in Face Bonus blocks, but only for fast trials (i.e. Q1 and Q2). For the model where motivation biased only the drift rate, the predicted proportion of scene responses in Scene Bonus blocks was higher than that in Face Bonus blocks across all four quartiles, with a relatively constant difference throughout. For the model where motivation biased both the starting point and drift rate, the predicted proportion of scene responses in Scene Bonus blocks was higher than that in Face Bonus blocks across all four quartiles, with the difference diminishing for slower trials. This pattern most closely resembles the pattern observed in participants' data. The CRT plots complement the linear regression and formal model comparison results to demonstrate how a model where motivation biased both the starting point and drift rate provided the best fit to participants' data. However, given the qualitative nature of the comparison and that it relies only on the trials at 48% scene, we have opted to keep the plots in the supplemental materials.

To assess how well a model where motivational bias in drift rate varied trial-by-trial with the evoked pupil response reproduced the relationship between evoked pupil response and motivationally biased categorizations, we simulated choice and response time data given the observed pupil dilation values and the posterior parameter estimates of model $v_p$. We averaged the evoked pupil response separately for trials on which the model predicted a motivation consistent response and for trials on which the model predicted a motivation inconsistent response, and performed a paired t-test to assess if mean evoked pupil response was different between the two trial types. The same analysis was then repeated with the empirical data for comparison (Fig. S7).

### Supplemental Figures

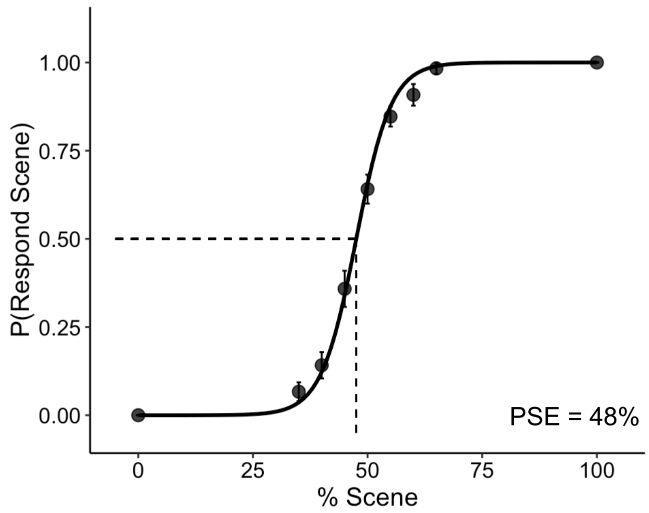


**Figure S1. Point of subjective equivalence (PSE) under neutral conditions.** Participants were equally likely to categorize an image as scene-dominant or face-dominant when the image comprised of 48% scene and 52% face.

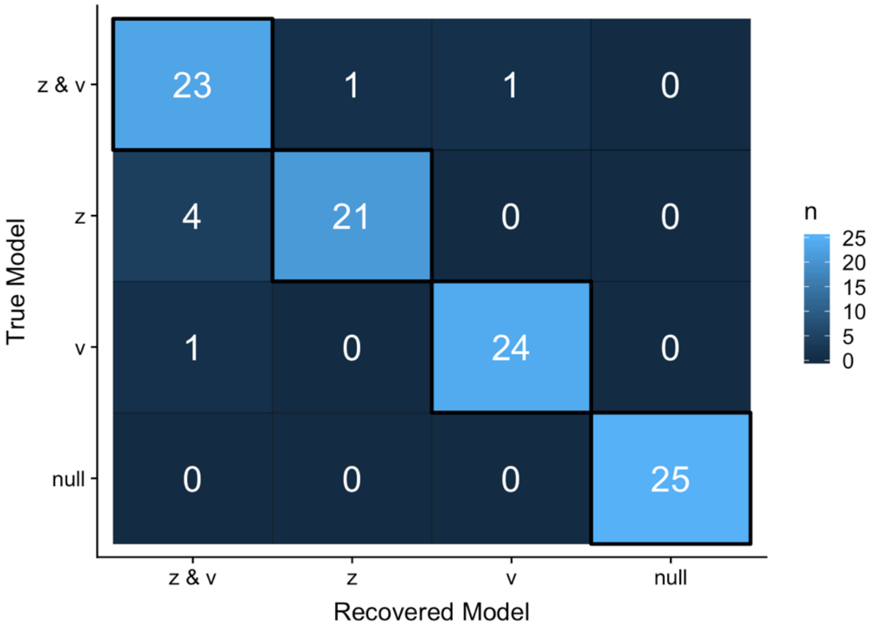


**Figure S2**. **Confusion matrix from model recovery simulations.** Each row indicates the number of simulated datasets (out of 25) on which data generated by a particular model is best fit by each of the models. The diagonal indicates accurate model recovery (e.g., correctly identifying *z & v* model when fitting to data generated by the *z & v* model).

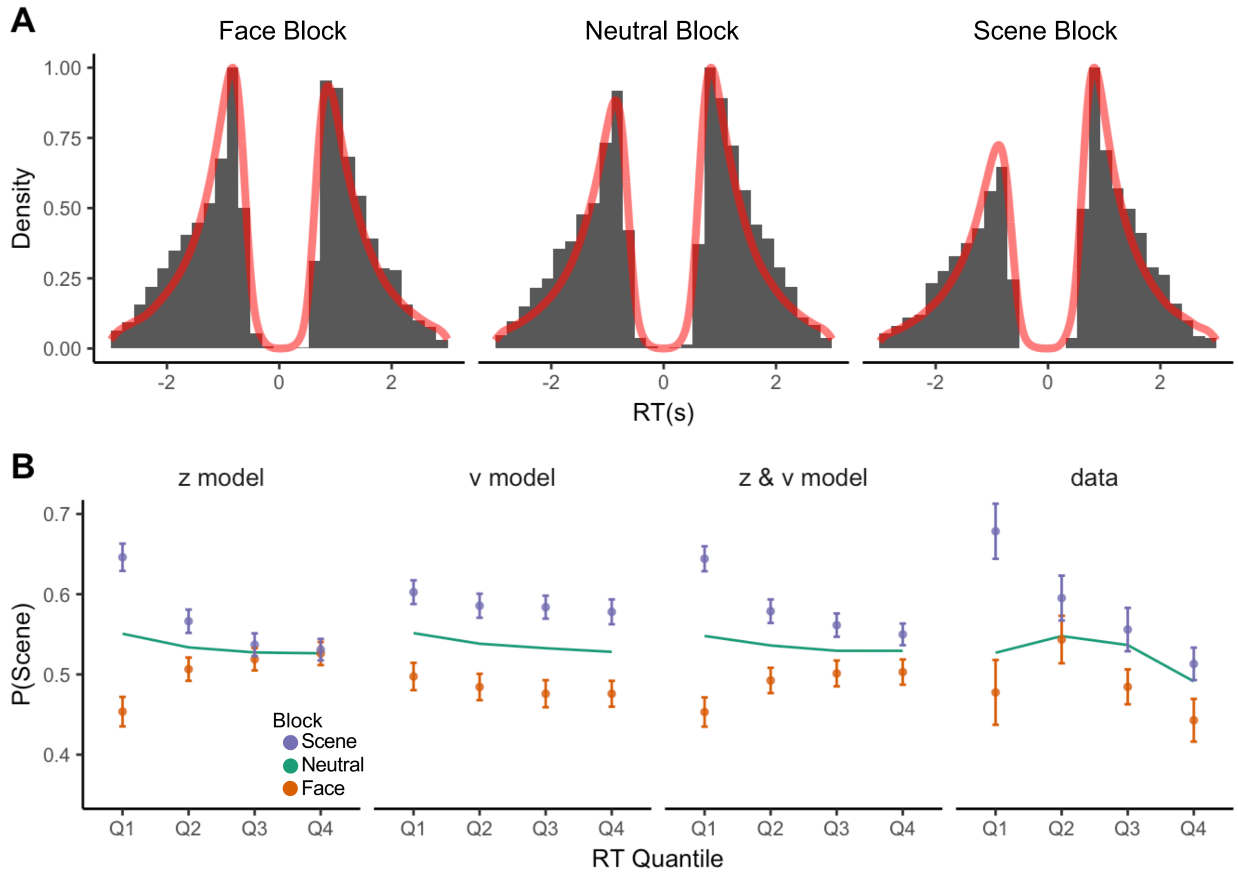


**Figure S3. Model where motivation biased both starting point and drift rate reproduced empirical pattern of response times. A. Observed and simulated response time distributions.** Histogram shows observed response time distributions separately for Face bonus, Neutral and Scene bonus blocks. Response time for face responses were sign-flipped for illustration purposes. Red line indicates simulated response time distributions from a model where motivation biased both the starting point and drift rate. **B. Conditional response time functions computed from observed and simulated data.** The response proportion was computed for each response time quartile and averaged across participants. In participants' data, the effect of motivation diminishes with longer response times, but was present even at the slowest quartile. This pattern was reproduced only in a model where motivation biased both the starting point and drift rate (*z & v* model, see Supplemental Results for additional discussion). Plot only includes trials at 48% scene as there were insufficient trials at the other levels of percent scene to be divided into quartiles. Error bars indicate between-subject standard error. To facilitate comparison between Face bonus and Scene bonus blocks, we plot the mean response proportions for the Neutral block as a line without error bars.

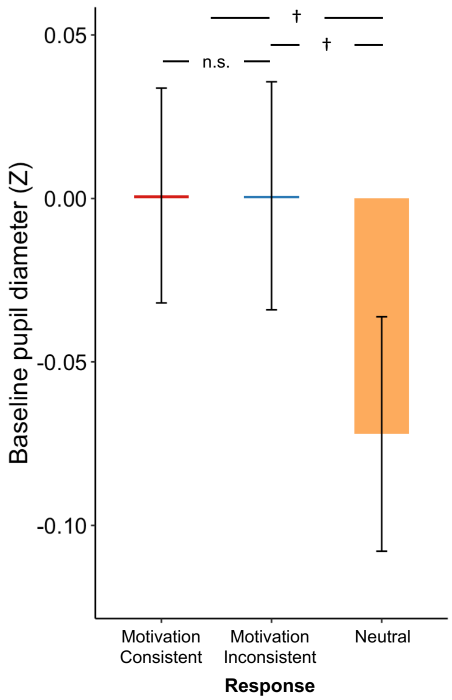


**Figure S4. Pre-stimulus baseline was not different prior to motivation consistent responses than motivation inconsistent responses but was marginally lower for neutral responses.** Average pupil dilation during the 500ms prior to stimulus onset separately for trials on which participants made motivation consistent responses (red), motivation inconsistent responses (blue) and neutral responses (i.e. trials in the neutral blocks; orange). Statistical significance was assessed using linear mixed effects models. n.s: not significant, †: $p < 0.10$.

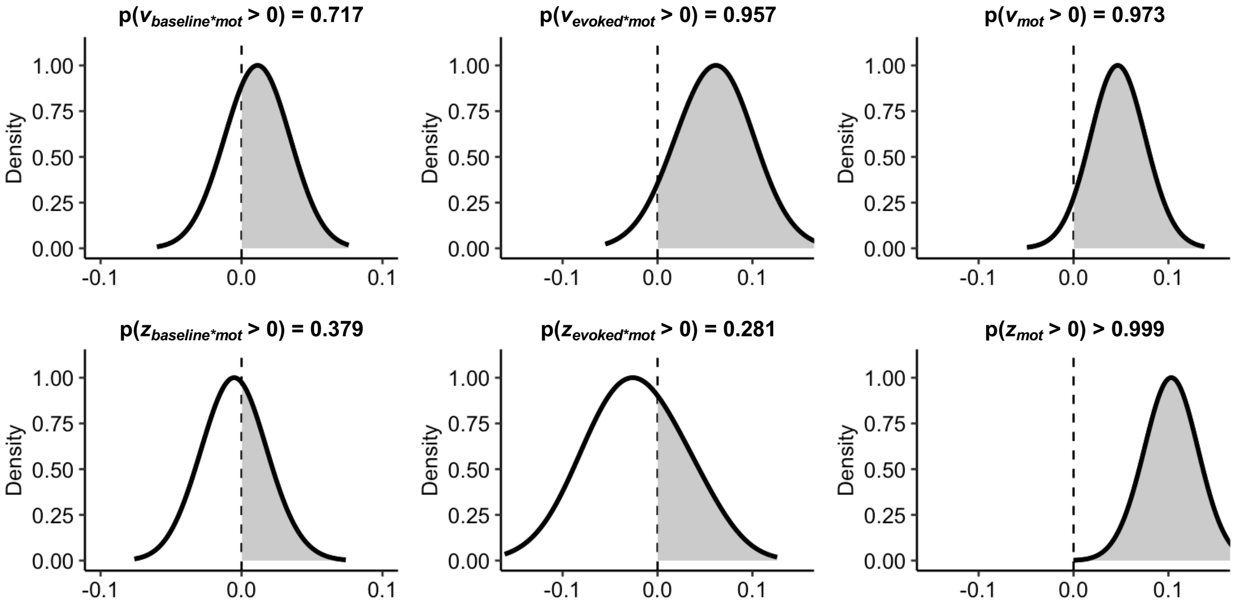


**Figure S5. Motivational biases in drift rate are associated with trial-by-trial fluctuations in evoked pupil response in model with the motivation consistent category as a covariate.** Posterior distributions for the regression coefficients of baseline pupil diameter on drift bias ($v_{baseline*mot}$) and starting point bias ($z_{baseline*mot}$), the regression coefficients of the evoked pupil response on drift bias ($v_{evoked*mot}$) and starting point bias ($z_{evoked*mot}$), the regression coefficients of the motivation consistent category on the drift rate ($v_{mot}$) and starting point ($v_{mot}$). Dashed line indicates 0 (no effect). There was a positive relationship between evoked pupil response and motivational biases in the drift rate ($p(v_{evoked*mot} > 0) > 0.95$)

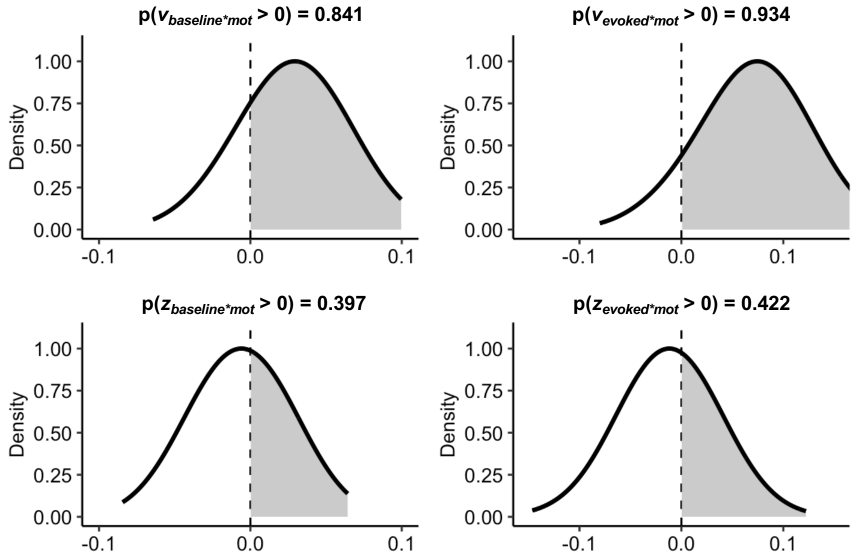


**Figure S6. Motivational biases in drift rate are associated with trial-by-trial fluctuations in evoked pupil response in a model with inter-trial variability parameters for drift rate, starting point and threshold.** Posterior distributions for the regression coefficients of baseline pupil diameter on drift bias ($v_{baseline*mot}$) and starting point bias ($z_{baseline*mot}$), and the regression coefficients of the evoked pupil response on drift bias ($v_{evoked*mot}$) and starting point bias ($z_{evoked*mot}$). Dashed line indicates 0 (no effect). There was a positive relationship between evoked pupil response and motivational biases in the drift rate, though the relationship fell short of a nominal p > 0.95 threshold ($p(v_{evoked*mot} > 0) = 0.934$), likely due to increased parameter uncertainty when including inter-trial variability parameters (Boehm et al., 2018).

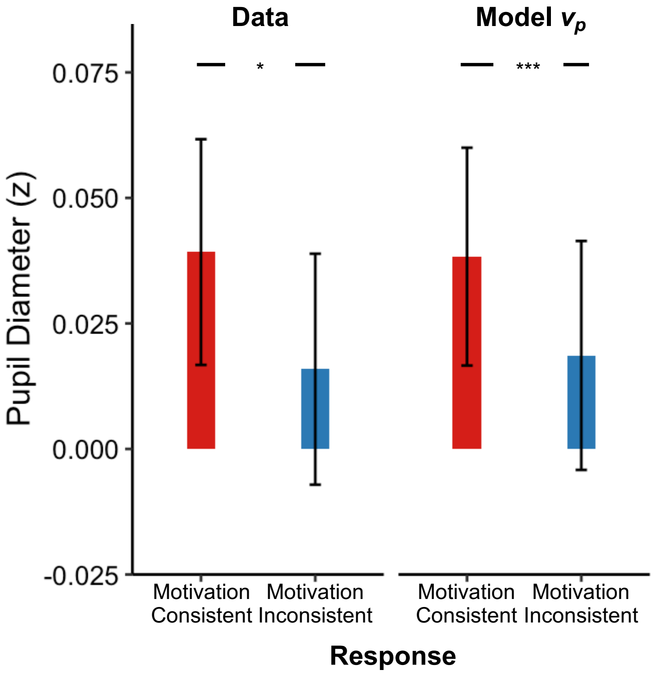


**Figure S7. Model $v_p$ reproduced the empirical observation that evoked pupil response was higher for motivation consistent responses than motivation inconsistent responses.** Evoked pupil response was average separately for trials on which the model predicted a motivation consistent response and for trials on which the model predicted a motivation inconsistent response. The analysis was then repeated with the empirical data for comparison. * $p < 0.05$, *** $p < 0.001$.

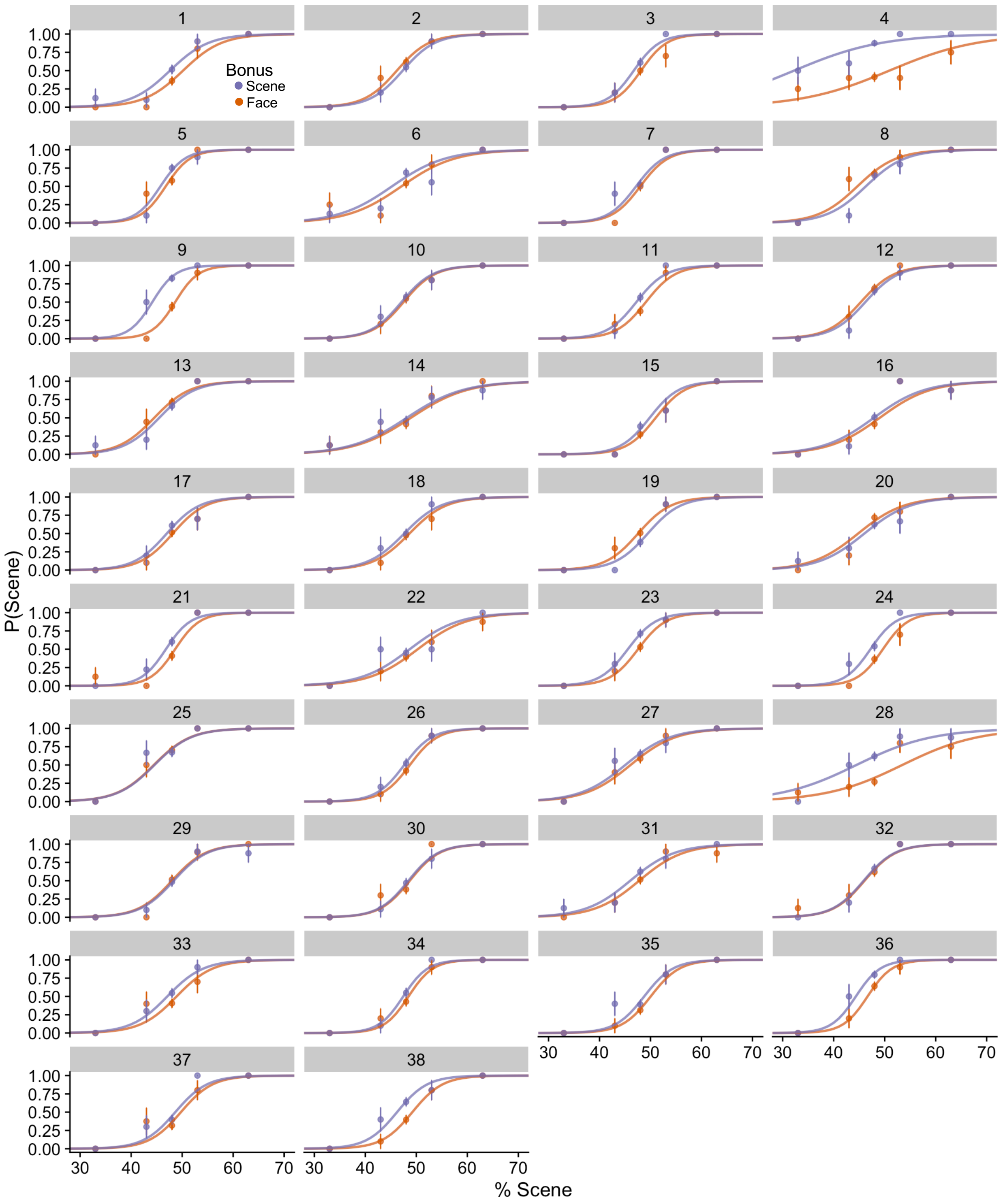


**Figure S8. Individual psychometric curves**. Proportion of scene-dominant responses as a function of percent scene, separately for Scene Bonus (purple) and Face Bonus (orange) blocks. Error bars indicate standard error of the mean. Trials from neutral blocks were excluded from the plots for visual clarity. Group-level differences between psychometric curves remain significant even when the three most extreme participants (#4, #9, #28) were excluded from the analyses ($b$ = 0.278, SE = 0.066, $z = 4.22$, $p < 0.001$)

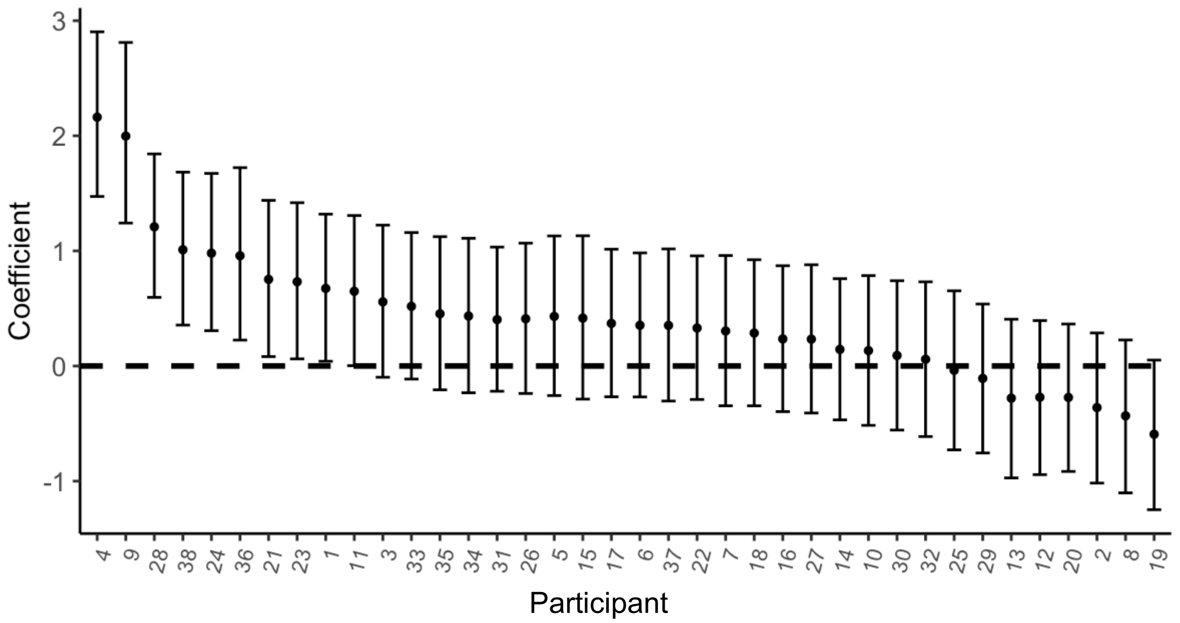


**Figure S9. Regression coefficients of effect of block type on the proportion of scene-dominant responses from fitting a GLM to each participant's data.** A positive value indicates higher proportion of scene-dominant responses on Scene Bonus blocks than Face Bonus blocks. Error bars indicate 95% confidence intervals.

### Supplemental Tables

**Supplementary Table 1**
**Model specification and estimated coefficients of linear mixed effects models.**

| | Formula | Term | Estimate | SE | *p* |
|---|---|---|---|---|---|
| **M1**[a] | response ~ % scene + condition + (% scene + condition \| subj) | intercept | -13.75 | 0.663 | 0.020 |
| | | % scene | 0.289 | 0.014 | < 0.001 |
| | | Face Bonus > Neutral | -0.145 | 0.070 | 0.037 |
| | | Scene Bonus > Neutral | 0.246 | 0.077 | 0.001 |
| **M2**[b] | RT ~ \|% scene - % face\| + response + response type + (\|% scene - % face\| + response + response type \| subj) | intercept | 1.445 | 0.048 | < 0.001 |
| | | \|% scene - % face\| | -0.304 | 0.019 | < 0.001 |
| | | response | -0.054 | 0.021 | 0.014 |
| | | Neutral > MotCon | 0.043 | 0.015 | 0.006 |
| | | MotIncon > MotCon | 0.031 | 0.012 | 0.018 |
| **M3**[c] | RT ~ \|% scene - % face\| + response + response type + (\|% scene - % face\| + response + response type \| subj) | intercept | 1.476 | 0.046 | < 0.001 |
| | | \|% scene - % face\| | -0.304 | 0.019 | < 0.001 |
| | | response | -0.054 | 0.021 | 0.014 |
| | | MotCon > Neutral | -0.031 | 0.012 | 0.010 |
| | | MotIncon > Neutral | 0.013 | 0.012 | 0.321 |
| **M4**[a,d] | response type ~ \|% scene - % face\| + baseline + evoked pupil + RT + (\|% scene - % face\| + baseline + evoked pupil + RT \| subj) | intercept | 0.477 | 0.123 | < 0.001 |
| | | \|% scene - % face\| | -0.190 | 0.057 | < 0.001 |
| | | baseline | 0.028 | 0.028 | 0.318 |
| | | evoked pupil | 0.166 | 0.057 | 0.004 |
| | | RT | -0.209 | 0.067 | 0.002 |
| **M5**[a] | accuracy ~ \|% scene - % face\| + baseline + evoked pupil + (\|% scene - % face\| + baseline + evoked pupil \| subj) | intercept | 0.055 | 0.140 | 0.694 |
| | | \|% scene - % face\| | 2.868 | 0.257 | < 0.001 |
| | | baseline | -0.040 | 0.060 | 0.506 |
| | | evoked pupil | -0.388 | 0.158 | 0.014 |
| **M6** | RT ~ \|% scene - % face\| + baseline + evoked pupil + (\|% scene - % face\| + baseline + evoked pupil \| subj) | intercept | 1.407 | 0.043 | < 0.001 |
| | | \|% scene - % face\| | -0.283 | 0.043 | < 0.001 |
| | | baseline | 0.018 | 0.013 | 0.172 |
| | | evoked pupil | 0.137 | 0.048 | 0.007 |

Note: Models are referred to by their labels (e.g., *M1*, *M2*, *M3*) in the *Methods* section. Formulas are written in the notation of the lme4 package in R with random effects indicated in parentheses. Variable coding - response: face = 0, scene = 1; condition: type of block, contrast coding with Neutral blocks as the reference group; RT = response time; response type: whether response was motivation consistent (MotCon), motivation inconsistent (MotIncon) or neutral; accuracy: whether the participant correctly categorized an image as face or scene-dominant. [a]A *logit* link function was used. [b]response type was coded with motivation consistent responses as the reference group. [c] response type was coded with neutral responses as the reference group. [d]neutral trials were dropped to allow for modeling using a GLMM.

**Supplementary Table 2**
**Model parameter estimates and convergence metrics**

| Parameters | Estimate | Gelman-Rubin $\hat{R}$ |
|---|---|---|
| ***Model without pupil data (Fig. 2A)*** | | |
| $a$ | 2.019 [1.926, 2.115] | 1.000 |
| $t$ | 0.488 [0.441, 0.541] | 1.000 |
| $z_{mot}$ | 0.100 [0.053, 0.145] | 1.001 |
| $z_{int}$ | 0.029 [0.017, 0.043] | 1.013 |
| $v_{mot}$ | 0.048 [0.000, 0.095] | 1.000 |
| $v_{\%scene}$ | 1.446 [1.325, 1.569] | 1.000 |
| $v_{int}$ | 0.059 [-0.007, 0.125] | 1.009 |
| ***Model with pupil data (Fig. 4A)*** | | |
| $a$ | 2.012 [1.919, 2.11] | 1.000 |
| $t$ | 0.486 [0.437, 0.538] | 1.000 |
| $z_{evoked*mot}$ | -0.003 [-0.082, 0.089] | 1.003 |
| $z_{baseline*mot}$ | -0.006 [-0.043, 0.031] | 1.004 |
| $z_{int}$ | 0.032 [0.022, 0.043] | 1.014 |
| $v_{evoked*mot}$ | 0.069 [0.004, 0.135] | 1.004 |
| $v_{baseline*mot}$ | 0.018 [-0.015, 0.058] | 1.004 |
| $v_{scene}$ | 1.427 [1.306, 1.55] | 1.000 |
| $v_{int}$ | 0.054 [-0.011, 0.12] | 1.003 |

Notes: Estimate denotes posterior mean of the corresponding parameter. Square brackets denote 95% credible interval. Convergence was assessed using the Gelman-Rubin $\hat{R}$ statistic. An $\hat{R}$ of greater than 1.1 would be indicative of nonconvergence. $t$: non-decision time, $z_{mot}$: effect of motivation on starting point, $z_{evoked*mot}$: effect of evoked pupil response on starting point bias, $z_{baseline*mot}$: effect of baseline pupil diameter on starting point bias, $z_{int}$: intercept of starting point, $v_{mot}$: effect of motivation on drift rate, $v_{evoked*mot}$: effect of evoked pupil response on drift bias, $v_{baseline*mot}$: effect of baseline pupil diameter on drift bias, $v_{\%scene}$: effect of percentage scene on drift rate, $v_{int}$: intercept of drift rate.

**Supplementary Table 3**
**Model comparison of models where motivational biases are modulated by pupil dilation**

| Model | DICc |
|---|---|
| $z_{bp}$_$v_{bp}$ | 26002 |
| $z_{bp}$_$v_p$ | 25997 |
| $z_b$_$v_{bp}$ | 25994 |
| $z_p$_$v_{bp}$ | 25998 |
| $z_b$_$v_p$ | 25991 |
| $z_p$_$v_p$ | 25986 |
| $v_{bp}$ | 25987 |
| $z_b$ | 25987 |
| $z_p$ | 25986 |
| $v_b$ | 25985 |
| $v_p$ | 25979 |
| *null* | 25986 |

Notes: Lower $DIC_c$ score indicates better model fit. *z* and *v* denote models where pupil dilation modulated starting point and drift bias respectively. Subscript *b* indicates that the parameter varied with baseline pupil diameter while subscript *p* indicates that the parameter varied with evoked pupil response. For example, $z_p$_$v_p$ indicates a model where motivational biases in both the starting point and drift rate varied with the evoked pupil response. *null* denotes a model where neither the starting point nor the drift rate varied with pupil dilation. A model where motivational biases in the drift rate varied with evoked pupil response (model $v_p$) provided the best fit to data.

**Supplementary Table 4**
**Participants' responses on post-experiment questionnaire**

| Sub # | Recoded Resp. | Raw Resp. (Were you influenced by the category bonuses?) |
|---|---|---|
| 1 | No | No |
| 2 | No | Not particularly |
| 3 | Yes | Yes |
| 4 | Sometimes | Sometimes |
| 5 | No | No |
| 6 | Yes | Yes |
| 7 | Sometimes | Yes, at times. |
| 8 | Sometimes | Sometimes. I tried to not be influenced by bonuses by focusing only on the gray screen in between photos and by reminding myself that picking the category of the bonus would not give me more money, only picking the right answer would. |
| 9 | Sometimes | I was influenced by the bonuses on the images I couldn't decide on |
| 10 | A little | Apart from the first few images per block, I think I forgot about them. |
| 11 | Yes | Yes |
| 12 | No | No |
| 13 | Yes | Yes |
| 14 | Yes | Yes, you were sometimes "hoping" to get a face/scene according to the category bonus. |
| 15 | Yes | Somehow yes |
| 16 | No | Not really. I just assumed I would get half of them and wouldn't get the other half. |
| 17 | No | No |
| 18 | Yes | Yes, so I tried to not read the name of the categories in order to not be biased |
| 19 | No | No |
| 20 | Yes | Yes |
| 21 | Yes | Yes |
| 22 | Yes | Yes |
| 23 | No | No |
| 24 | No | No |
| 25 | No | No |
| 26 | No | No |
| 27 | A little | Slightly |
| 28 | A little | Loosely |
| 29 | A little | To a very little extent |
| 30 | Yes | Yes |
| 31 | Yes | Yes |
| 32 | No | No |
| 33 | Yes | Yes |
| 34 | Yes | Yes |
| 35 | A little | Only a little at times |
| 36 | Yes | Yes |
| 37 | No | Not really, since my answers do not change whether or not I got paid. |
| 38 | Yes | Yes |
